# Effect of the p75 Neurotrophin Receptor on Glycemic Control of Obese Mice during Time-Restricted Feeding

**DOI:** 10.1101/2020.08.19.257394

**Authors:** Brandon Podyma, Katherine Battin, Dove-Anna Johnson, Ali D. Güler, Christopher D. Deppmann

## Abstract

Circulating glucose regulates organismal energy homeostasis and is tightly controlled in response to feeding behavior. In overweight individuals, glucose homeostasis is often perturbed due to resistance to normal satiety signals, such as insulin and leptin, leading to type 2 diabetes and its attendant complications. An emerging dietary intervention, time-restricted feeding (TRF), aims to ameliorate the adverse metabolic consequences of obesity, however, its effectiveness on glucose control is uncertain. Here, we demonstrate that TRF is only transiently effective in reducing pre-meal serum glucose levels in obese mice lacking leptin. However, in *Ob/Ob* mice that also lack a gene known to suppress behavior associated with TRF, the p75 neurotrophin receptor (p75NTR), a sustained reduction of glucose levels is observed. These results suggest that the effectiveness of TRF on glycemic control can be enhanced with concurrent targeting of modulators of glucose homeostasis and TRF response.

## INTRODUCTION

Metabolic health is increasingly appreciated as a fundamental requirement in preventing many chronic diseases (1). Obesity and type 2 diabetes come with symptoms that are highly systemic, and confer an increased risk for further complications such as cardiovascular disease and cancer (2). Compounding their detrimental effects is the lack of a broadly effective means to reverse obesity or type 2 diabetes (3).

Recently a novel diet regimen, time-restricted feeding (TRF), has gained attention as a plausible option for improving metabolic health by leveraging our knowledge of circadian biology to better time food intake to match organismal metabolic activity (4). Benefits of TRF in several clinical studies of patients at risk for obesity and type 2 diabetes have included increased insulin sensitivity and lowering of blood pressure, with several other trials ongoing in those with obesity and the metabolic syndrome (5–7). However, while TRF has shown some improvement of glucose homeostasis in mice on a high fat diet, the effect in humans is less clear, with only modest benefits observed (4–6,8). As glucose control is one of the most important drivers of detrimental health outcomes in those with obesity and diabetes, this suggests that combinatorial strategies may be necessary to achieve optimal health benefits from TRF (9).

Animal models of obesity have historically been used to study the control of glucose homeostasis in the context of increased weight and adiposity. One of the best characterized genetic models of obesity is the leptin-deficient *Ob*/*Ob* mouse, which develops massive obesity due to increased food intake and decreased energy expenditure, and subsequent hyperglycemia (10,11). Notably, the *Ob*/*Ob* background has been used previously to identify that the soluble ligand TNFα and its receptors (TNFR1, *Tnfrsf1*; TNFR2, *Tnfrsf2*) are necessary to promote hyperglycemia and insulin resistance in obesity (12,13). More recently we found a novel signaling interaction between TNFα/TNFR signaling with a paralogous TNFR superfamily member, the p75 neurotrophin receptor (p75NTR, *Ngfr*) (14,15). Given these connections, we questioned whether there may be a similar role for p75NTR in obesity.

Similar to the TNFα receptors, p75NTR has been implicated in insulin sensitivity and glucose control in adipocytes, and has also been found to be necessary to promote weight gain on a high fat diet (16,17). Recently, we found that mice lacking p75NTR (*Ngfr*-KO) exhibit a phenotype of decreased food intake with increased weight loss on TRF, but while maintaining blood glucose levels similar to that of controls (18). However, due to their resistance to diet-induced obesity, it is unknown whether p75NTR may have a role in modulating glucose homeostasis during TRF in the setting of obesity.

To study this possibility, we generated a genetic model of *p75NTR* deletion on a background of leptin-deficient obesity (*ob/ob*). We used this model to test the hypothesis that targeting p75NTR function during TRF would lead to improvements in glycaemic control despite obesity. Unexpectedly, we find that this dual loss of leptin and p75NTR leads to a variable weight phenotype, with increased mortality in adolescent and young adult mice due to an unknown cause. In mice that reach full adulthood, however, we find strikingly improved glucose homeostasis independent of body weight during TRF.

## RESULTS

### Loss of p75NTR in leptin-deficient *ob/ob* mice leads to obesity and hyperglycemia with increased mortality

To examine the physiologic effect of p75NTR loss in leptin-deficient mice, body weight and food intake were measured over the course of 15 weeks. Male *Ob*-*Ngfr*KO mice and WT controls had similar body weights until around week 7. From week 7 to week 15, approximately 40% of *Ob*-*Ngfr*KO mice began losing weight (**Fig 1A**). This weight loss occurred concurrently with decreased food intake, and all cases of weight loss in *Ob*-*Ngfr*KO mice progressed to the point of death or removal from the study by 15 weeks of age (**Fig 1B, C**). In those *Ob*-*Ngfr*KO mice that survived, we find they gained weight similarly to controls, with a similar ratio of feed efficiency despite a lower level of absolute food intake (**Fig 1D-F**). While the discordance between body weight and food intake may suggest a possible change in energy output, as has been previously demonstrated for *Ngfr*KO mice (17), it remains unknown whether *Ob*-*Ngfr*KO mice have decreased energy expenditure. We also demonstrate that baseline blood glucose is similar between genotypes (**Fig 1G**).

**Figure 1.**
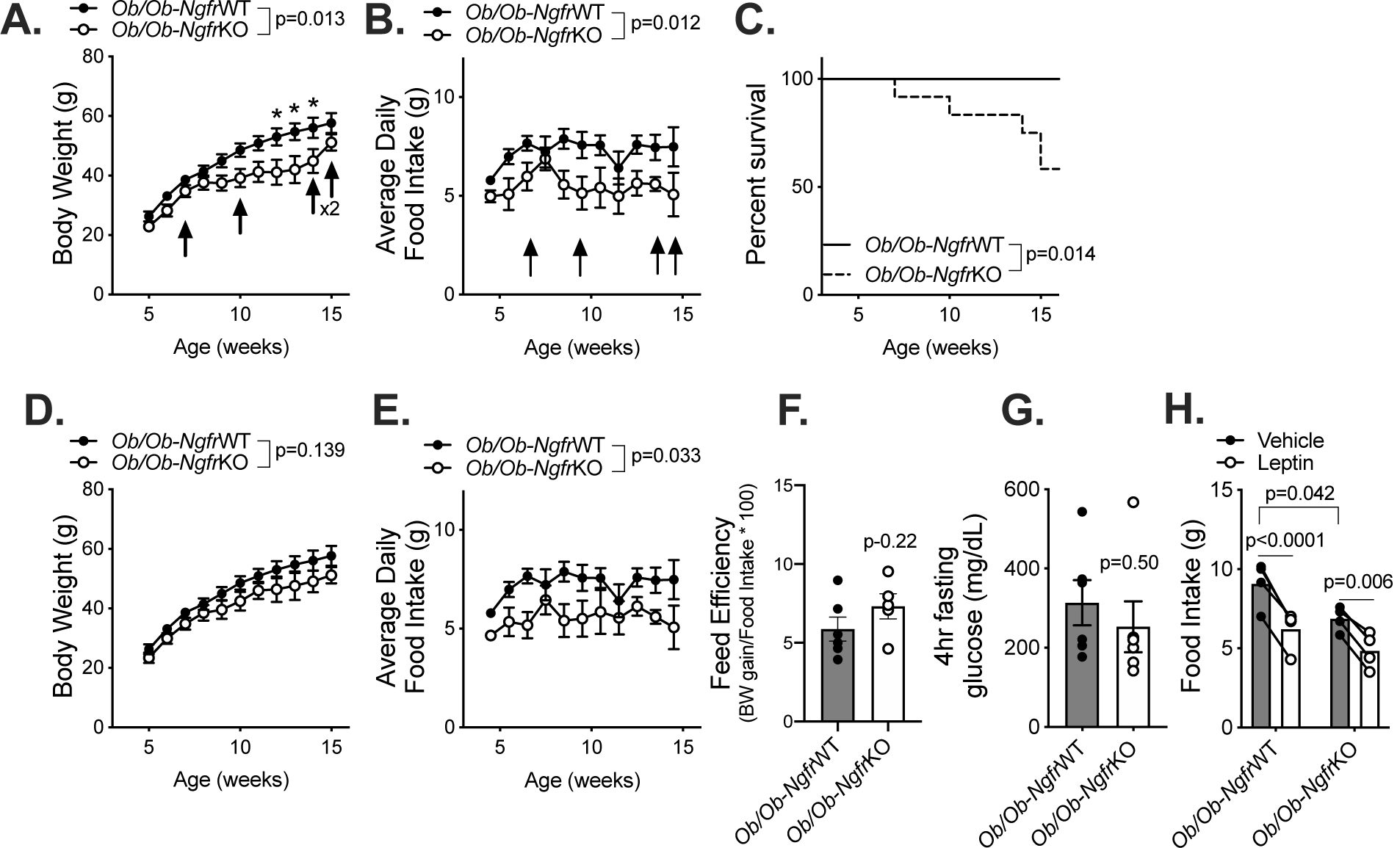
*Ob*-*Ngfr*KO *mice develop obesity and hyperglycemia, but with increased mortality*. (**A**) Male mouse body weight curves during adolescence into adulthood, n=12/group. Arrows indicate the death of an *Ob*-*Ngfr*KO mouse. F(1,22)=7.256, p=0.013 overall between genotypes by mixed effects Anova. *p<0.01 at time points between genotypes by mixed effects Anova with Bonferroni’s multiple comparisons post-hoc test. (**B**) Daily food intake averaged over the course of 7 days during development, n=7 *Ob*-*Ngfr*WT, 8 *Ob*-*Ngfr*KO. Arrows indicate the death of an *Ob*-*Ngfr*KO mouse. F(1,13)= 8.446, p=0.012 between genotypes by mixed effects Anova. (**C**) Survival curve during development, n=12/group. p=0.014 by Mantel-Cox test (χ^2^=6.06, df=1). (**D**) Male mouse body weight curves of surviving mice from adolescence into adulthood (replotted from A), n=12 WT, 7 KO. F(1,17)=2.408, p=0.139 between genotypes by two-way Anova. (**E**) Daily food intake of surviving mice averaged over the course of 7 days during development (replotted from B), n=7 *Ob*-*Ngfr*WT, 5 *Ob*-*Ngfr*KO. F(1,10)= 6.131, p=0.033 between genotypes by two-way Anova. (**F**) Feed efficiency (10 week body weight gain (g) divided by 10 week food intake (g)) of survivors. n=7 *Ob*-*Ngfr*WT, 5 *Ob*-*Ngfr*KO. F(1,10)= 6.131, p=0.22 between genotypes by Student’s t-test. (**G**) Blood glucose following a 4 hour daytime fast in 12 week old mice, n=6/group. p=0.50 by Student’s t-test. (**H**) Food intake over 24 hours following i.p. vehicle or leptin (1 ug) administration just before lights out, n=4/group. F(1,6)=4.79 between genotypes by two-way Anova with Bonferroni’s multiple comparisons test.

As p75NTR has been implicated in other metabolic functions regulated by leptin, including feeding and glucose homeostasis, we next assessed whether leptin responsiveness was intact (16,18). Following intraperitoneal vehicle administration, *Ob*-*Ngfr*KO mice ate significantly less than controls (**Fig 1G**), agreeing with the prior trend in food intake data (**Fig 1B, 1E**). Upon leptin administration (1ug, i.p.), however, both genotypes exhibited similar reductions in food intake (**Fig 1G**).

### Blood glucose levels are improved in *Ob*-*Ngfr*KO mice during time restricted feeding

To examine the effect of p75NTR on glucose homeostasis in the setting of obesity, *Ob-Ngfr*WT and surviving *Ob-Ngfr*KO mice were placed on a five day TRF (**Fig 2**). While *Ob-Ngfr*KO mice weighed around 9% less than controls, we observed similar trends in body weight and food intake over the course of restriction, with no statistically significant differences at any time point (**Fig 2A, B**). Both genotypes also experienced a similar reduction in blood glucose during the initial fast, however, blood glucose levels returned to normal in *Ob-Ngfr*WT mice by day 3 while they remained significantly lower in *Ob-Ngfr*KO mice **(Fig 2C)**. While loss of p75NTR alone does not significantly alter fed or fasting insulin levels, *Ngfr*KO mice have been demonstrated to have improved insulin sensitivity (16,18). However, it is unknown whether *Ob-Ngfr*KO mice have similarly improved insulin sensitivity, nor whether this could explain the observed changes in glucose. Thus, it appears that *Ob-Ngfr*KO mice on TRF are able to adaptively correct their hyperglycemia in response to food restriction, while *Ob*-*Ngfr*WT mice are not.

**Figure 2.**
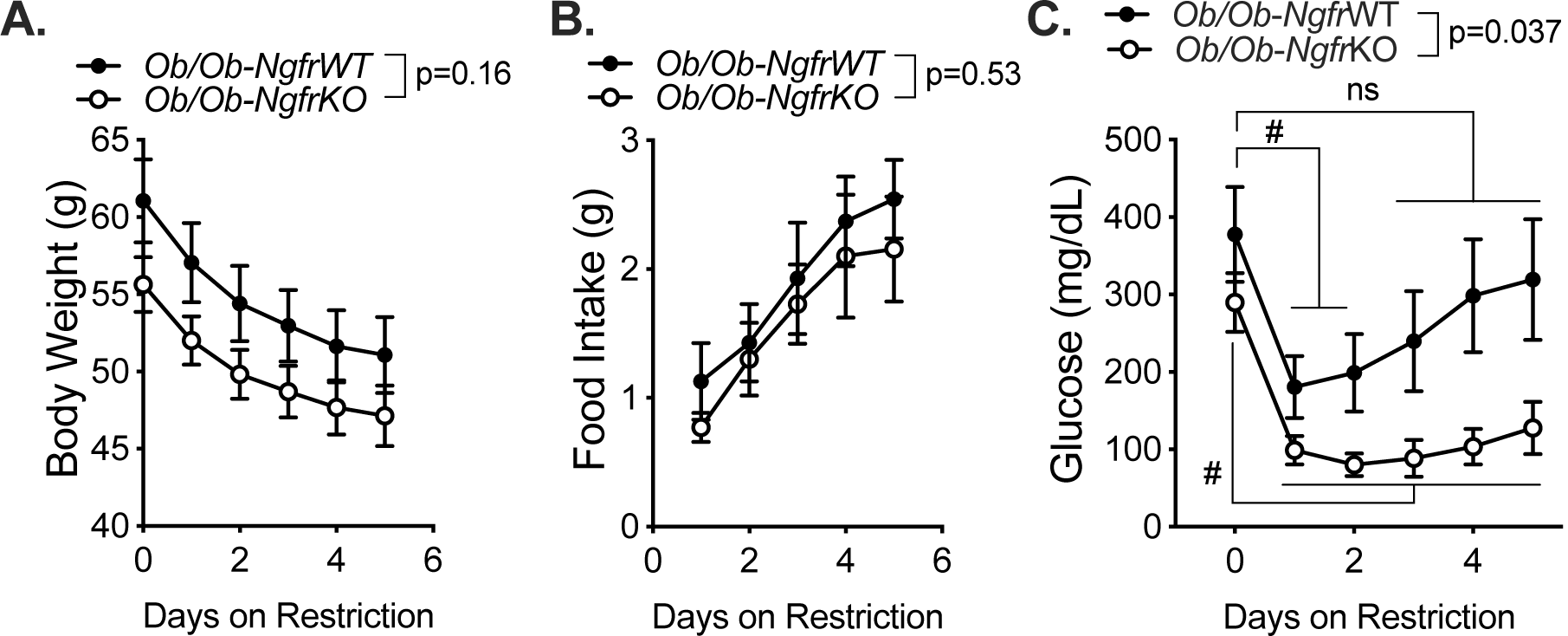
*Ob*-*Ngfr*KO *mice have significantly lower blood glucose during TRF compared to controls*. 16-20 week old mice were placed onto TRF, whereby food was removed just before lights off on day 0, and was made available for only three hours per day on each of the next five days. Mice were allowed *ad libitum* access to food during the 3 hour meal time, and body weight and blood glucose measurements were made daily just after lights on. (**A**) Body weight of mice on TRF. F(1,12)=2.29, p=0.16 between genotypes by two-way Anova. (**B**) *Ad libitum* food intake during 3 hour mealtime. F(1,12)=0.428, p=0.53 between genotypes by two-way Anova. (**C**) Blood glucose measurements of mice on TRF. F(1,12)=5.48, p=0.037 between genotypes by two-way Anova. #p<0.05 compared to day 0 blood glucose by two-way Anova with Bonferroni’s multiple comparisons.

## DISCUSSION

Time restricted feeding is a metabolic intervention with increasing evidence for wide ranging benefits, though with mixed support for improvements in glycemic control. Pairing TRF with a targeted therapy against molecular players involved in glucose homeostasis and the response to TRF could be an effective strategy to further improve metabolic outcomes. Here, we demonstrate that obese mice with a loss of the p75NTR have similar baseline body weight and hyperglycemia as controls, but exhibit a larger and longer lasting reduction in blood glucose on TRF. A major limitation of this study, however, is the survival heterogeneity of mice with a double knockout of leptin and p75NTR. Elucidating the cause of their loss of body weight and appetite could provide a new window on our understanding of leptin and neurotrophin signaling in energy balance.

Previous work has delineated a role for p75NTR in reducing insulin sensitivity by decreasing Rab5 activity and increasing Rab31 activity, thereby reducing GLUT4 in the plasma membrane (16). In *Ngfr*KO mice fed a normal chow diet, this leads to an increase of glucose uptake in adipocytes and myocytes, decreasing overall serum glucose levels (16). It was unknown, however, whether these improvements would be maintained in the setting of increased weight, as *Ngfr*KO mice are resistant to diet-induced obesity. We suggest here that these improvements are indeed maintained during obesity, and that this provides preliminary evidence that targeting p75NTR function could improve the efficacy of TRF regimens with the aim of improving glycemic control. However, the approximate 9% baseline difference in body weight observed limits this interpretation, and warrants further investigation using strategies of weight matched cohorts. Further, given the unexpected lethality of some of our *Ob*-*Ngfr*KO mice, further study is warranted to establish the cause of death and possible adverse side effects in the acute setting, especially with the use of continuous glucose monitoring for possible drop-out effects.

Achieving improved glucose control is a clinically important endpoint for individuals with obesity and type 2 diabetes, and has been shown to significantly improve health outcomes. We believe coupling TRF with strategies to improve glucose homeostasis provides a promising path forward to improving metabolic health.

## METHODS

### Mice

All experiments were carried out in compliance with the Association for Assessment of Laboratory Animal Care policies and approved by the University of Virginia Animal Care and Use Committee. Animals were housed on a 12-h light/dark cycle with food (Teklad Diet 8664) and water *ad libitum* unless otherwise indicated. *Ngfr-KO* mice were purchased from Jackson Labs (Bar Harbor, ME, USA) (#002213)(19), and were maintained on a B6;129s mixed background and genotyped with primers against intron II in *Ngfr -* Intron II (Ngfr-IntII, 5′-CGA TGC TCC TAT GGC TAC TA), Intron III (*Ngfr*-IntIII, 5′-CCT CGC ATT CGG CGT CAG CC), and the pGK-Neo cassette (pGK, 5′-GGG AAC TTC CTG ACT AGG GG). *Ob/Ob* mice were purchased from Jackson Labs (#000632) and genotyped by PCR amplifying (Forward primer: TGT CCA AGA TGG ACC AGA CTC, Reverse primer: ACT GGT CTG AGG CAG GGA GCA) a region around the Ob point mutation, followed by restriction digest using Dde1. All crosses were carried out using *Ob*/+ and *Ngfr*+/- or *Ngfr*-/- mice. All data shown is for male mice.

### Body Weight and Food Intake

Food intake was performed on individually housed male mice that were acclimated for 7 days, followed by food and body weight measurements weekly (for development curves), or daily during restricted feeding experiments.

### Leptin Administration

Vehicle (100 ul saline) or recombinant leptin (1 ug in 100 ul saline; Peprotech, Rocky Hill, NJ, USA) was injected intraperitoneally into 12-16 week old *ad libitum* fed mice just prior to lights out. Food intake was measured 24 hours later.

### Time Restricted Feeding

For scheduled feeding, mice were first acclimated to single housing for 7 days. For daytime scheduled feeding, mice were fasted at lights off (ZT12) on day 0. Mice were weighed, and glucose (one touch ultra 2, Bayer, Leverkusen, Germany) measures were taken 12 hours later at lights on (ZT0). Mice were then refed 4 hours later at ZT4, with food removed 3 hours later at ZT7. Mice were fed between ZT4-7 on each of the next four days.

### Statistical Analysis

Data are presented as mean ± SEM. Statistical analysis was carried out using Prism version 8.0 (Graphpad, San Diego, CA, USA). Student’s t-test was used to compare single means between genotypes, and 2-way Anova was used to compare genotype by time interactions. Differences were considered significant if *p* < 0.05.

## ACKNOWLEDGEMENTS

We would like to thank members of the Deppmann and Güler labs for helpful feedback during all stages of this work. This work was supported by T32-GM7267-39 and T32-GM7055-45 to BP, Hartwell Foundation to CDD, and R01-GM121937 to ADG.

